# Imputing missing distances in molecular phylogenetics

**DOI:** 10.1101/276345

**Authors:** Xuhua Xia

## Abstract

Missing data are frequently encountered in molecular phylogenetics and need to be imputed. For a distance matrix with missing distances, the least-squares approach is often used for imputing the missing values. Here I develop a method, similar to the expectation-maximization algorithm, to impute multiple missing distance in a distance matrix. I show that, for inferring the best tree and missing distances, the minimum evolution criterion is not as desirable as the least-squares criterion. I also discuss the problem involving cases where the missing values cannot be uniquely determined, e.g., when a missing distance involve two sister taxa. The new method has the advantage over the existing one in that it does not assume a molecular clock. I have implemented the function in DAMBE software which is freely available at available at http://dambe.bio.uottawa.ca

## 1 Introduction

The demand to reconstruct supertrees with a large number of taxa from concatenating a large number of genes often results in missing data illustrated in Fig. 1 where a distance between Sp3 and Sp4 cannot be computed because they share no homologous sites. While missing data can be accommodated by the likelihood method with the pruning algorithm [1, 2, 3, pp. 253-255], they can inflate branch lengths and introduce phylogenetic bias [4, 5]. Some popular likelihood-based phylogenetic methods, e.g., PhyML [6], use distance-based methods to build the initial phylogenetic tree, which is then modified in various ways and evaluated in the likelihood framework to find the maximum likelihood tree. Distance-based methods are much faster than other phylogenetic methods such as maximum likelihood, Bayesian inference and maximum parsimony. Consequently, they are used frequently constructing supertrees [7].

**Fig. 1.**
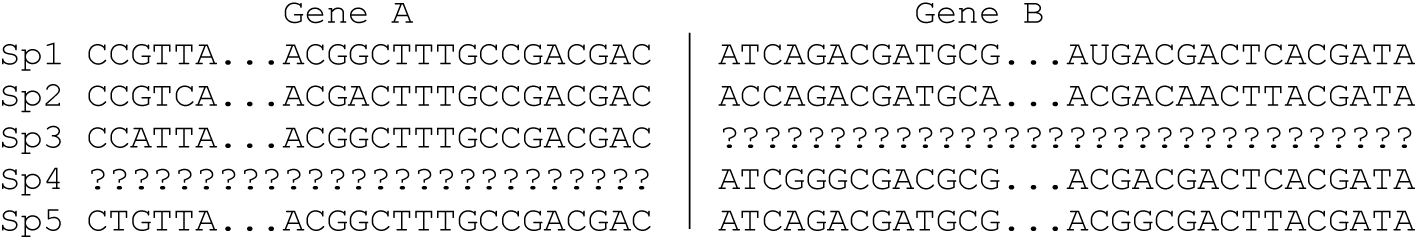
A sequence data set from concatenating Gene A and Gene B sequences. A distance cannot be computed between Sp3 and Sp4 because they share no homologous sites.

The least-squares (LS) method is frequently used for imputing the missing distances. The general conceptual framework is to estimate the missing distances given different trees, and then choose the best tree and the best estimates of missing distances with a specific criterion. One of its key advantages of the LS method lies in its optimization with constraints that restrict branch length to be non-negative. I will first outline the general approach, point out problematic cases where unique solution cannot be found, and then develop an efficient computational method similar to the expectation-maximization (EM) algorithm to impute the missing distance. I illustrate the method by applying it to real data.

## 2 Least-square method for imputing missing distances

Suppose we have four species (S1 to S4 in Fig. 2) with D_12_ = 2, D_14_ = 5, D_23_ = 3, D_24_ = 5, D_34_ =4 but with D_13_ missing. The method developed can also be used to impute multiple missing distances.

**Fig. 2.**
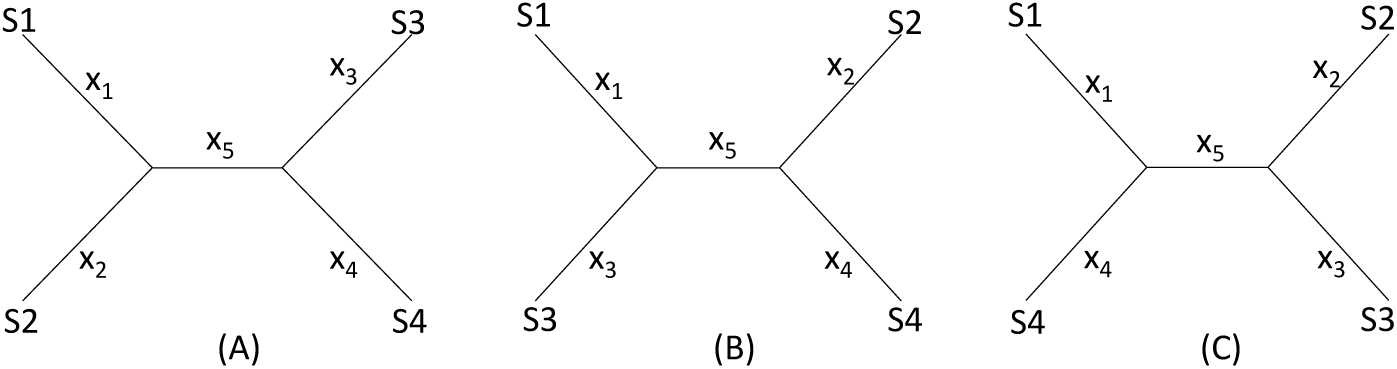
Topologies for illustrating the distance-based methods

### 2.1 A wrong approach

One may take a wrong approach by thinking that, in this particular case, we have five unknowns and five equations and can solve for D_13_ exactly. For example, given a topology in Fig. 2A, we can write the expected D_ij_ values, i.e., E(D_ij_), as:

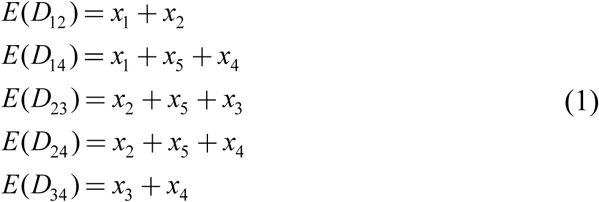

These E(D_ij_) values are termed patristic distances in phylogenetics. If we replace E(D_ij_) by the observed D_ij_ values, we can indeed solve the simultaneous equations in Eq. (1), which give the solution as

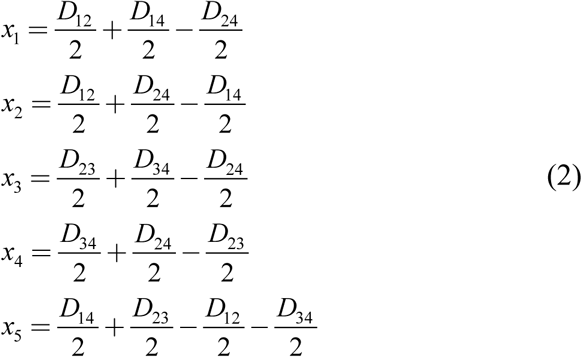

The missing D_13_ given the tree in Fig. 2A, designated as D_13.A_, can therefore be inferred, as:

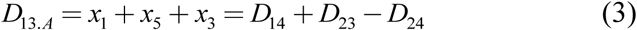

Thus, given the five known D_ij_ values above, we can obtain *x*_*1*_ = *x*_*2*_ = *x*_*3*_ = *x*_*5*_ = 1, *x*_*4*_ = 3, D_13.A_ = 3. The tree length (TL), defined as TL = ∑*x*_*i*_, is 7 for the tree in Fig. 2A, i.e., TL_A_ = 7.

One might think of applying the same approach to the other two trees in Fig. 2B,C to obtain D_13.B_ and D_13.C_ as well as TL_B_ and TL_C_, and choose as the best D_13_ and the best tree by using the minimum evolution criterion [8, 9], i.e., the tree with the shortest TL.

This approach has two problems. First, the approach fails with the tree in Fig. 2B where the missing distance, D_13_, involves two sister species. One can still write down five simultaneous equations, but will find no solutions for *x*_*i*_, given the D_ij_ values above, because the determinant of the coefficient matrix is 0. For the tree is Fig. 2C, the solution will have x_5_ = -1. A negative branch length is biologically undesirable and defeats the ME criterion for choosing the best tree and the associated estimate of D_13_. Second, in most practical cases where missing distances are imputed, there are more equations than unknowns, e.g., if we have five or more species with one missing distance.

### 2.2 Least-squares approach

Take E(D_ij_) specifications in Eq. (1) for the tree in Fig. 2A, the LS approach find D_13_ and the best tree that minimize the residual sum of squared deviation (RSS) between the observed D_ij_ and the expected E(D_ij_):

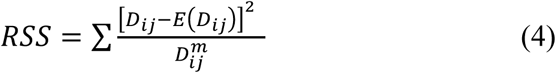

where m is typically 0 (ordinary least-squares, OLS), 1, or 2. In the illustration below, we will take the OLS approach with m = 0. It has been shown before that OLS actually exhibits less topological bias than alternatives with m equal to 1 or 2 [10]

Given the three tree topology, the results from the LS estimation are summarized in Table 1. Note that, for the tree in Fig. 2B, there are multiple sets of solutions of *x*_*i*_ that can achieve the minimum RSS of 1.

**Table 1.** Estimation results from minimizing RSS, with Trees A, B, and C as in Fig. 2, and with the constraint of no negative branch lengths.

| Site | Tree A | Tree B | Tree C |
| --- | --- | --- | --- |
| $x_1$ | 1 | 0 | 1 |
| $x_2$ | 1 | 1.5 | 1.5 |
| $x_3$ | 1 | 0 | 1 |
| $x_4$ | 3 | 3.5 | 3.5 |
| $x_5$ | 1 | 1 | 0 |
| $D_{13}$ | 3 | 0 | 2 |
| TL | 7 | 6 | 7 |
| RSS | 0 | 1 | 1 |

We see a conflict between the LS criterion and the ME criterion in choosing the best tree and the best estimate of D_13_. The ME criterion would have chosen Tree B with TL_B_ = 6 and D_13_ = 0 because TL_B_ is the smallest of the three TL values. In contrast, the LS criterion would have chosen Tree A with RSS = 0 and D_13_ = 3. There is no strong statistical rationale for the ME criterion which is based on the assumption that substitutions are typically rare in evolution. The ME criterion is particularly inappropriate for imputing missing distances because it tends to underestimate the missing distances. In contrast, the LS-criterion is well-established. I recommend the LS criterion for the simultaneous imputing of missing distances and inferring phylogenetic trees. Phylogeneticists sometimes think that the ME criterion would be appropriate if the branch lengths are not allowed to take negative values [11–13]. The illustrative example in Table 1 shows that, even when we do not allow branch lengths to become negative, there is still problem with the ME criterion.

## 3 Implementation in DAMBE

An earlier version of DAMBE implemented the LS approach above by using an iterative approach similar to the EM (expectation-maximization) algorithm as follows. For a given distance matrix with missing values, we simply fill in the missing D_ij_ values by guestimates, e.g., the average of the observed distances. These initial D_ij_ guestimates will be designated as D_ij,m0_ where the subscript “m0” indicates missing distances at step 0. We now build a tree from the distance matrix that minimizes RSS in Eq. (4). From the resulting tree we obtain the patristic distances E(D_ij_) from the tree and replace D_ij.m0_ by the corresponding E(D_ij_) values which are now designated as D_ij.m1_. We now again build a tree, obtain the corresponding E(D_ij_) to replace D_ij.m1_ so now we have D_ij.m2_. We repeat this process until RSS does not decrease any further. This process can quickly arrive at a local minimum. Unfortunately, different topologies have different minimums, and this approach is too often locked in a local minimum with a tree that does not achieve a global minimum RSS.

New version of DAMBE (since version 7) uses a downhill simplex method in multidimensions {Press, 1992 #26508, pp. 408-412} multiple times (with different initial values for the points in the simplex) to increase the chance of finding the global RSS associated with the missing distances and the tree. When there is a single missing distance, then the Brent’s method {Press, 1992 #26508, pp. 402-408} is used. Fig. 3 shows an illustrative example. The distance matrix in Fig. 3a is computed from aligned sequence data used before [14]. Fig. 3b is the phylogenetic tree built from this distance matrix. Suppose D_gibbon,orangutan_ and D_gorrila.chimpazee_ are missing (shaded in Fig. 3a) and need to be imputed. DAMBE yields D_gibbon,orangutan_ = 1.3776 and D_gorrila.chimpazee_ = 0.4600, which are close to the observed values (Fig. 3a). The final tree built from the distance matrix with the two missing distances is identical to Fig. 3b except for a negligible difference in branch lengths.

**Fig. 3.**
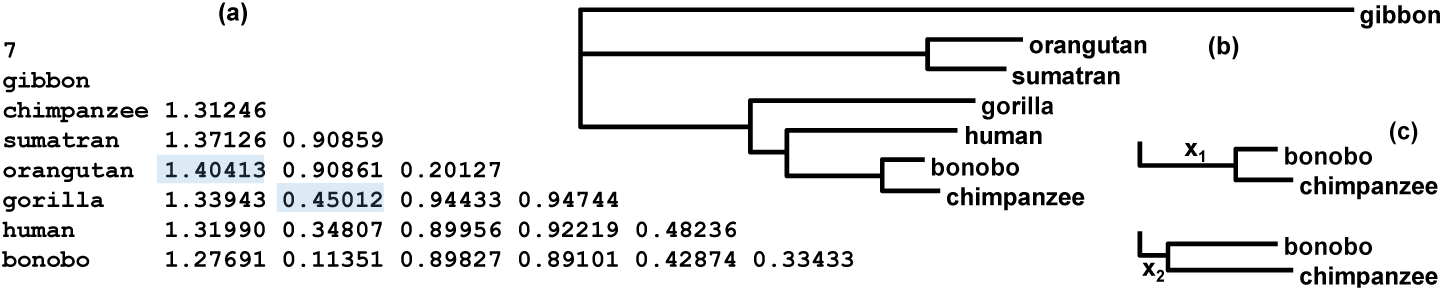
An example data set for imputing missing distances. (a) A real distance matrix computed from aligned sequences, but we pretend that the two shaded distances are missing. (b) A phylogenetic tree from the distance matrix. (c) A special case illustrating the problem of the estimation.

One can access the function in DAMBE [15, 16] by clicking ‘File|Open other molecular data|Distance matrix file with missing value’, and open a distance file in the format in Fig. 3a with missing distances represented by ‘.’ (a period without quotation marks).

There are cases where missing distances cannot be uniquely determined. For example, when the missing distance is for two sister taxa (e.g., the two chimpanzee species, designated bonobo and chimpanzee in Fig. 3b), we can find a minimum RSS but the solution for the missing distance D_bonobo,chimpanzee_ is not unique. That is, multiple D_bonobo,chimpanzee_ values can generate the same RSS. Note that the patristic distances E(D_bonobo.i_) and E(D_chimpanzee.i_) where i stands for other species, do not depend on the branch length leading to the common ancestor from bonobo and chimpanzee. This branch can be as short as x_2_ or as long as x_1_ (Fig. 3c) but RSS in Eq. **Error! Reference source not found**. will remain the same. Thus, in this particular case, a missing D_bonobo,chimpanzee_ cannot be determined uniquely. The only way to eliminate this problem is to have a more closely related species to break up the sister relationship so that the missing D_ij_ is not between two sister taxa.

The method in the paper has an advantage over a previous method [7] that assumes a rooted tree and a molecular clock for building a tree and for inferring missing distances. This assumption is not needed and is too restrictive in practise.

## 4 Software availability

DAMBE is available free at http://dambe.bio.uottawa.ca. One can access the function by clicking ‘File|Open other molecular data|Distance matrix file with missing value’, and open a distance matrix file in the format in Fig. 3a, with missing distances represented by ‘.’ (a period without quotation marks).

## 5 Acknowledgement

This study is funded by the Discovery Grant from Natural Science and Engineering Research Council (NSERC, RGPIN/261252) of Canada.

## 6 Conflict of interest

I declare no competing interest.

